# Diffusion Tensor Imaging: Influence of segmentation on fiber tracking in the supraspinatus muscle – an inter-operator reliability analysis

**DOI:** 10.1101/2023.05.15.540752

**Authors:** Sebastian Vetter, Hans-Peter Köhler, Pierre Hepp, Hanno Steinke, Stefan Schleifenbaum, Jan Theopold, Simon Kiem, Maren Witt, Jeanette Henkelmann, Christian Roth

**Affiliations:** Department of Biomechanics in Sports, Leipzig University, Leipzig, Germany; Department of Orthopedics, Trauma and Plastic Surgery, Universitätsklinikum, Leipzig, Germany; Department of Anatomy, Universitätsklinikum, Leipzig, Germany; ZESBO - Zentrum zur Erforschung der Stütz- und Bewegungsorgane, Universitätsklinikum, Leipzig, Germany; Institute of Sport and Motion Science, University of Stuttgart, Stuttgart, Germany; Department of Diagnostic and Interventional Radiology, Universitätsklinikum, Leipzig, Germany; Department of Paediatric Radiology, Universitätsklinikum, Leipzig, Germany

## Abstract

The ability of muscle to generate force depends on its architecture and quality. MR-based diffusion tensor imaging of muscle (mDTI) is an innovative approach for showing the fiber arrangement for the whole muscle volume. For accurate calculations of fiber metrics, muscle segmentation prior to tractography seems mandatory. Since segmentation is known to be operator dependent, it is important to understand how segmentation affects tractography. The aim of this study was to compare the results of deterministic fiber tracking based on muscle models generated by two independent operators. In addition, this study compares the results with a segmentation-free approach. Fifteen subjects underwent mDTI of the right shoulder. The results showed that mDTI can be successfully applied to complex joints such as the human shoulder. Furthermore, operators segmentation did not influence the results of fiber tracking and showed excellent intraclass correlation estimates (≥ 0.975). As an exploratory approach, the segmentation-free fiber tracking showed significant differences in terms of mean fascicle length. Based on these findings, we assume that tractography does not require a detailed muscle segmentation. Furthermore, it implies that mDTI and automatic segmentation approaches or even a segmentation-free analysis can be considered for evaluation of muscle changes.

## Introduction

Muscle architecture is a primary determinant of its function. A dramatic change in muscle fiber arrangement occurs with changes in sarcomeres[1–3]. The addition of sarcomeres in series lengthens the muscle fascicle[4], increases the shortening velocity[5], develops the range of motion[6] and supports force buffering and energy absorption during decelerative motion[7]. In practice, a longer fascicle length (FL) is positively correlated with sport-specific performance[8–11]. Consequently, knee flexor weakness and a shorter FL have been shown to increase the risk of injury[12–15]. Therefore, FL is a primary parameter to determine the muscle architecture and the potential force output of the muscle. Since science and practice have paid much attention to the volume and cross-sectional area of a muscle, research into whole muscle architecture and quality[16,17] has long been underrepresented[18] and undervalued[19].

To investigate muscle architecture, the most common non-invasive method is ultrasound imaging. Ultrasound appears to be a convenient, robust, and relatively inexpensive imaging technique[20]. However, ultrasound has several disadvantages: limited field of view, poor representation of all muscle fibers, difficult and time consuming to standardise[21]. This leads to difficulties in using ultrasound to access changes in muscle fiber arrangement over time[22,23]. The fact that a change in muscle architecture is not uniform across all regions of the muscle[24], shows that an accurate method and a macroscopic view of a muscle’s fiber arrangement is at least as important as a method’s economics. Therefore, an innovative technology such as DTI may be of interest. DTI is an already accepted and valid method in neuroscience and shows the arrangement of nerves by measuring the diffusion of water molecules along the nerve fibers in the brain[25]. As muscles also contain a certain amount of water, an mDTI also provides a valid[26–28], reliable and robust[29,30] calculation of muscle fiber metrics. In addition, this tool can be used to examine even more metrics that also describe muscle quality[16,17,31]. Nevertheless, mDTI also has certain drawbacks: measurement of surrogate biomarkers, reconstruction of tracts varies and depends on algorithms and analysis methods, MR field strength, vendor and sequences[32] and is time-consuming to process data[20,33].

In order to turn mDTI into a tool of practical significance, it is worth taking a look at the most common data processing steps. Among several processing methods commonly used in this field[34–36], the slice-by-slice segmentation of muscle seems to be the most laborious and operator-dependent step[37,38]. Segmentation is necessary to generate a region of interest (ROI) for each subject[38]. Since it is known that ROI-based tractography does not stop at the surface of the ROI[30] and that different segmentation techniques alter the tractography result for the lower limb muscles[37], the question is how different segmentation routines affect mDTI analysis for the human shoulder. Furthermore, it seems interesting to check the importance of segmentation for an accelerated mDTI approach. Therefore, this study aims to show the dependence of two different segmentation routines for segmentation-based analysis (SBA 1 and SBA 2) on the manually segmented muscle model volume (MV) and its influence on the deterministic fiber tracking result. Furthermore, in an exploratory way, we want to compare these results with a segmentation or model-free approach (MFA). For the reliability analysis, we focused on the calculation of muscle shape indices (MV, FL and fiber volume [FV]) and conventional reported tensor metrics (fractional anisotropy [FA], axial diffusivity [AD], radial diffusivity [RD] and mean diffusivity [MD]), which reflect the arrangement and quality of a muscle[16,17]. As ultrasound is a conventional and convenient tool to determine the fiber architecture, we aim to discuss ultrasound and mDTI in terms of feasibility in research and daily clinical routine.

## Materials and Methods

This study analysed a dataset from an interventional trial. The protocol was approved by the local ethics committee (Leipzig University, No: 362/21-ek) and is registered in the German Register for Clinical Trials (DRKS00028885). The study was conducted in accordance with the relevant guidelines and regulations. Written informed consent was obtained from all subjects. The authors had no access to information that could identify individual participants during or after data collection.

### Sample

Fifteen healthy male subjects (24.1 ± 3.8 years of age; 183.7 ± 6.5 cm tall; 79.5 ± 7 kg body mass) were included in this study. Criteria for inclusion were: no history of muscle-nerve disease, no injuries, no discomfort or pain in the right shoulder, no resistance training five days prior to data collection, no regular medication.

### Data acquisition

Magnet resonance imaging (MRI) of the right shoulder was performed using a 3 Tesla MRI scanner (Siemens MAGNETOM Prisma Fit, Erlangen, Germany) with a dedicated 16-channel shoulder coil in the head-first supine position. The right shoulder was placed in a neutral position with the arm adducted and the hand supinated. The MR protocol consisted of a 3D coronal T1-weighted (T1w) and a sagittal DTI sequence from distal to proximal. The total scan time was approximately twelve minutes. The T1w sequence was acquired with the following parameters: repetition and echo time *TR/TE* = 492/20 ms, slice thickness = 0.7 mm, flip angle = 120°, field of view FOV = 180 × 180 mm^2^, matrix = 256 × 256 mm^2^. For DTI a commercial Siemens 2D echo planar diffusion image sequence was acquired with the following parameters: repetition and echo time *TR/TE* = 6100/69 ms, slice thickness = 4 mm, flip angle = 90°, field of view FOV = 240 × 240 mm^2^, matrix = 122 × 122 mm^2^, 48 diffusion sampling directions with *b* = 400 s/mm^2^.

### Muscle segmentation

Manual segmentation was based on common mDTI methods described previously. Segmentation was performed using Mimics Materialise (v.24.0, Leuven, Belgium). Two independent operators (SBA 1 and SBA 2) segmented each M. supraspinatus. Segmentation was based on the recorded T1w sequence of each subject. To compare individual differences in segmentation, each operator generated an individual segmentation routine for the whole data set. The first segmentation step was to generate a base mask by setting a threshold on the grey values of the images to separate muscle-tendons from bony structures. Then, both operators split the basic muscle mask to separate the M. supraspinatus from the other surrounding tissues and to proceed with manual segmentation and correction. While SBA 1 preferred manual segmentation, operator two (SBA 2) focused on interpolation using the integrated multi-slice editing function (Figure 1). However, both operators used semi-automatic segmentation functions and differed in time spent on each step. Finally, each surface model was smoothed by a factor of 0.5 and exported as a ROI for fiber tracking.

**Figure 1.**
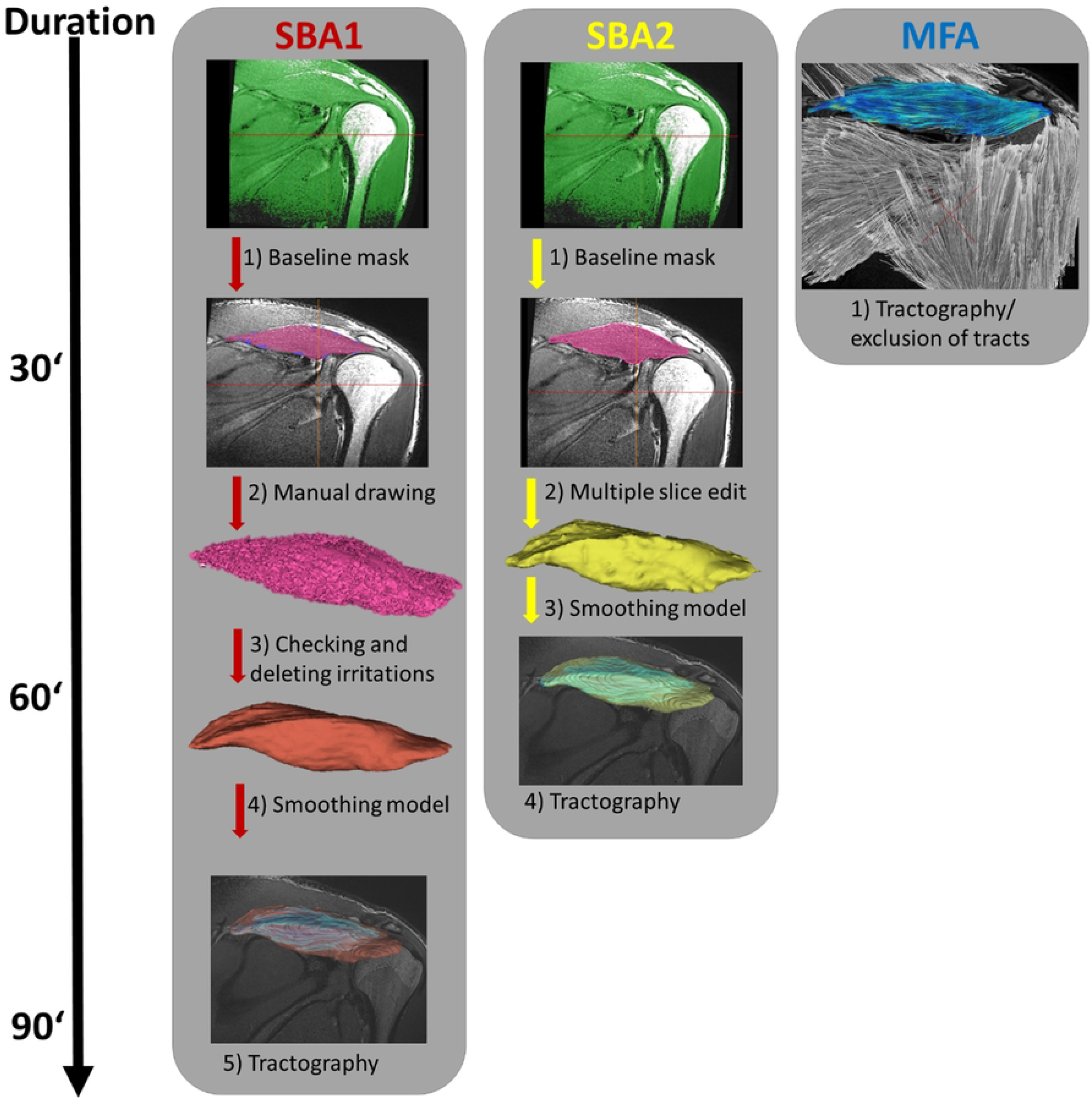
Workflow of methods. Workflow displays different processing steps and each methods duration in minutes (‘). Segmentation-based analysis by operator 1 (SBA 1) included four major segmentation steps, operator 2 (SBA 2) displayed three steps. Model-free analysis (MFA) did not include a segmentation and used the entire field of view as seeding area for deterministic fiber tracking. Within MFA the red cross symbolises the manual exclusion of tracts outside of the highlighted M. supraspinatus (blue color).

### DTI data processing and fiber tracking

DSI Studio (v. 3^th^ of December 2021. http://dsi-studio.labsolver.org) was used for DTI processing, deterministic fiber tracking and tract calculations. To perform tractography for the M. supraspinatus, we registered and resampled the DTI images to the T1w images. The quality of the DTI and FA maps was first visually checked by two experts using DSI Studio. In addition, the DTI images were corrected for motion and eddy current distortion using DSI Studio’s integrated FSL eddy current correction. To ensure plausible fiber tracking results, we used the following stopping criteria recommended: maximum angle between tract segments 15°, 20 mm ≤ tract length ≤ 130 mm; step size = 1.5 mm. These settings were oriented to FL results of cadaveric dissections[39] and recommendations for deterministic muscle fiber tracking stopping criteria[40–42].

Fiber tracking was then performed either within a model ROI (SBA methods) or for the entire DTI images without using a segmented model (MFA). After a reconstruction of ∼10.000 tracts for the M. supraspinatus region, tractography was terminated and duplicates were deleted. Since MFA used the entire DTI image as a seeding area for tractography we removed all tracts outside the M. supraspinatus. Next, clearly implausible tracts and tracts crossing the muscle boundary within the SBA and MFA were reviewed and removed by two experts. Finally, DTI tensor parameters (FA, AD, MD and RD) and muscle parameters (MV, FL and FV) were calculated based on a deterministic fiber tracking algorithm[43] and specific tracking strategies[31] using DSI Studio. Since MFA did not include a muscle segmentation step, it took approximately 30 minutes. In contrast, SBA 1 and SBA 2, including segmentation, took approximately 90 and 60 minutes respectively.

### Data analysis

SPSS v.27 (IBM. Armonk. New York. USA) was used for statistics. Figures were generated using MATLAB v.R2022a (MathWorks. Natick. USA). Descriptive results were based on the calculation of mean values and the standard deviation (±). As the human shoulder has complex structures and very different muscle shapes, two raters visually inspected the image slices for quality. If the inspection or statistics revealed outliers, they were excluded from further analysis. Outliers were defined visually based on distribution and box plots and calculation of z-transformed values. If they exceeded 3.0 within a metric, they were excluded. After checking the quality of the data, repeated measures analysis of variance (rmANOVA) was used to show overall differences in means between methods. Greenhouse-Geisser correction was applied if appropriate. When a main effect was found, Bonferroni-corrected post-hoc comparisons were performed. In addition, interrater-reliability between all methods (SBA 1, SBA 2 and MFA) was analysed using an intraclass correlation coefficient (ICC) for each mDTI (FA, MD, AD, RD) and muscle (FL, FV, and MV) parameter. ICC estimates and their 95%-confidence intervals (CI) were based on a single-rating, absolute-agreement and a two-way random effects-model (2.2). ICC values less than 0.5 were considered as poor, 0.5 to 0.75 as moderate, 0.75 to 0.9 as good, and greater than 0.9 as excellent[44]. The significance level was set at *p* < .05. Bland-Altman plots were calculated to show the limits of agreement.

## Results

All MR scans could be used for data processing and were included in the analysis. Segmentation for SBA 1 and SBA 2 was performed by two independent operators. SBA 1, SBA 2 and MFA were applied to the human M. supraspinatus of the right shoulder.

The descriptive results (Table 1) show similar values with low variance for mDTI indices (CV ≤ 8.02). In contrast, FL and FV show higher variance (CV ≥ 19.45) and mean differences between subjects and methods (FL mean = 36.34 – 41.27 mm). Greenhouse Geyser corrected rmANOVA were used to assess differences and showed a main model effect only for FL (F1.135, 15.890 = 22.645; *p* < 0.001; η^2^_p_ = 0.618). Further post-hoc paired *t*-tests (Figure 2) revealed significant differences in segmentations with respect to MV (t14 = 6.407; *p* = 0.001), but no significant differences in FL between the two SBA methods (Figure 2).

**Table 1.** Descriptive results.

| Metric | SBA 1 |  |  |  |  | SBA 2 |  |  |  |  | MFA |  |  |  |  |
| --- | --- | --- | --- | --- | --- | --- | --- | --- | --- | --- | --- | --- | --- | --- | --- |
| | Mean | ± | Min. | Max. | CV % | Mean | ± | Min. | Max. | CV % | Mean | ± | Min. | Max. | CV % |
| FA | 0.28 | 0.02 | 0.25 | 0.32 | 7.76 | 0.28 | 0.02 | 0.25 | 0.32 | 8.02 | 0.28 | 0.02 | 0.25 | 0.31 | 7.38 |
| MD | 1.87 | 0.06 | 1.78 | 2.00 | 3.34 | 1.87 | 0.06 | 1.77 | 1.99 | 3.35 | 1.87 | 0.07 | 1.79 | 1.99 | 3.59 |
| AD | 2.44 | 0.09 | 2.27 | 2.64 | 3.62 | 2.44 | 0.09 | 2.28 | 2.63 | 3.56 | 2.45 | 0.09 | 2.30 | 2.61 | 3.70 |
| RD | 1.58 | 0.06 | 1.47 | 1.68 | 3.81 | 1.58 | 0.06 | 1.46 | 1.67 | 3.90 | 1.59 | 0.07 | 1.49 | 1.68 | 4.10 |
| FL | 36.34 | 8.12 | 26.59 | 49.31 | 22.35 | 36.41 | 8.16 | 26.27 | 48.09 | 22.40 | 41.27 | 8.03 | 29.60 | 55.37 | 19.45 |
| FV | 137.6 | 31.3 | 87.8 | 179.3 | 22.8 | 133.8 | 32.5 | 84.5 | 177.5 | 24.3 | 139.6 | 30.4 | 80.7 | 191.7 | 21.8 |

Columns show the mean value, standard deviation ( ± ), minimum and maximum values and coefficient of variation (CV %) for each method, segmentation-based analysis (SBA) and model-free analysis (MFA). The rows show fractional anisotropy (FA), axial diffusivity (AD), radial diffusivity (RD), and mean diffusivity (MD) in $10^{-3}$ mm/s, fascicle length (FL) in mm, and fiber volume (FV) in $\text{mm}^3$ .

**Figure 2.**
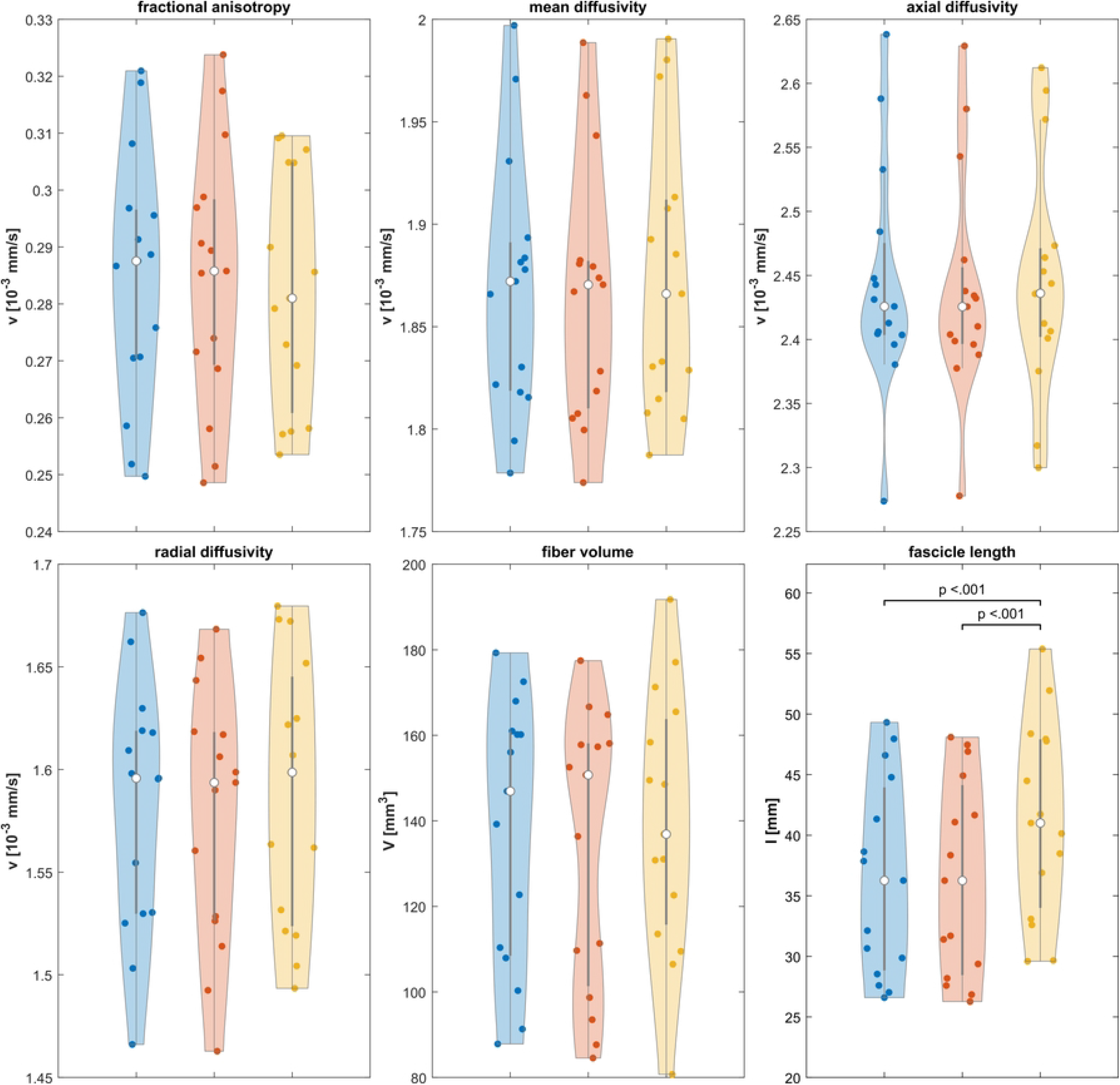
Violin plots and mean differences between methods. Violin plots show mean differences between segmentation-based analysis (SBA 1 [blue] and SBA 2 [red]) and model-free analysis (MFA, yellow).

When assessing operator dependence for fiber tracking outcome measures (Table 2) the ICC was found to be good (≥ 0.751). Agreement was highest between SBA 1 and SBA 2, ranging from ICC estimates between 0.975 to 0.997. As an explorative approach, the comparison of SBA to MFA revealed an ICC of 0.751 for FL. Ignoring the results for FL, the ICC values for MFA varied between 0.840 (FV) and 0.930 (MD).

**Table 2.** Interrater reliability.

| Parameter | Methods | ICC | 95% CI |  |
| --- | --- | --- | --- | --- |
| FA | Overall | 0.935 | 0.856 | 0.976 |
|  | SBA 1 vs SBA 2 | 0.997 | 0.992 | 0.999 |
|  | SBA 1 vs. MFA | 0.901 | 0.736 | 0.965 |
|  | SBA 2 vs. MFA | 0.903 | 0.741 | 0.966 |
| MD | Overall | 0.950 | 0.887 | 0.981 |
|  | SBA 1 vs SBA 2 | 0.994 | 0.979 | 0.998 |
|  | SBA 1 vs. MFA | 0.928 | 0.804 | 0.975 |
|  | SBA 2 vs. MFA | 0.930 | 0.808 | 0.976 |
| AD | Overall | 0.948 | 0.883 | 0.981 |
|  | SBA 1 vs SBA 2 | 0.995 | 0.980 | 0.999 |
|  | SBA 1 vs. MFA | 0.923 | 0.787 | 0.974 |
|  | SBA 2 vs. MFA | 0.927 | 0.801 | 0.975 |
| RD | Overall | 0.944 | 0.875 | 0.979 |
|  | SBA 1 vs SBA 2 | 0.994 | 0.981 | 0.998 |
|  | SBA 1 vs. MFA | 0.920 | 0.786 | 0.972 |
|  | SBA 2 vs. MFA | 0.923 | 0.783 | 0.973 |
| FL | Overall | 0.823 | 0.401 | 0.944 |
|  | SBA 1 vs SBA 2 | 0.990 | 0.972 | 0.997 |
|  | SBA 1 vs. MFA | 0.751 | 0.012 | 0.933 |
|  | SBA 2 vs. MFA | 0.752 | 0.000 | 0.932 |
| FV | Overall | 0.888 | 0.761 | 0.957 |
|  | SBA 1 vs SBA 2 | 0.975 | 0.908 | 0.992 |
|  | SBA 1 vs. MFA | 0.840 | 0.588 | 0.943 |
|  | SBA 2 vs. MFA | 0.843 | 0.604 | 0.944 |

Intraclass coefficient (ICC) and its confidence intervals (CI) for segmentation-based analysis (SBA 1 and SBA 2) and model-free analysis (MFA) for the parameters fractional anisotropy (FA), axial diffusivity (AD), radial diffusivity (RD), and mean diffusivity (MD) in $10^{-3}$ mm/s, fascicle length (FL) in mm, and fiber volume (FV) in mm<sup>3</sup>.

Bland-Altman plots (Figure 3) display the mean and the difference of each method within the limits of agreement. The lowest limits of agreement were found for SBA 1 and SBA 2, with the exception of FV(3.78 ± 12.20). The highest limits of agreement show comparisons between SBA and MFA similar for FL (4.87 and 4.93 for SBA 2 vs MFA) but different for the metric FV (1.98 and 5.76 for SBA 2 vs MFA).

**Figure 3.**
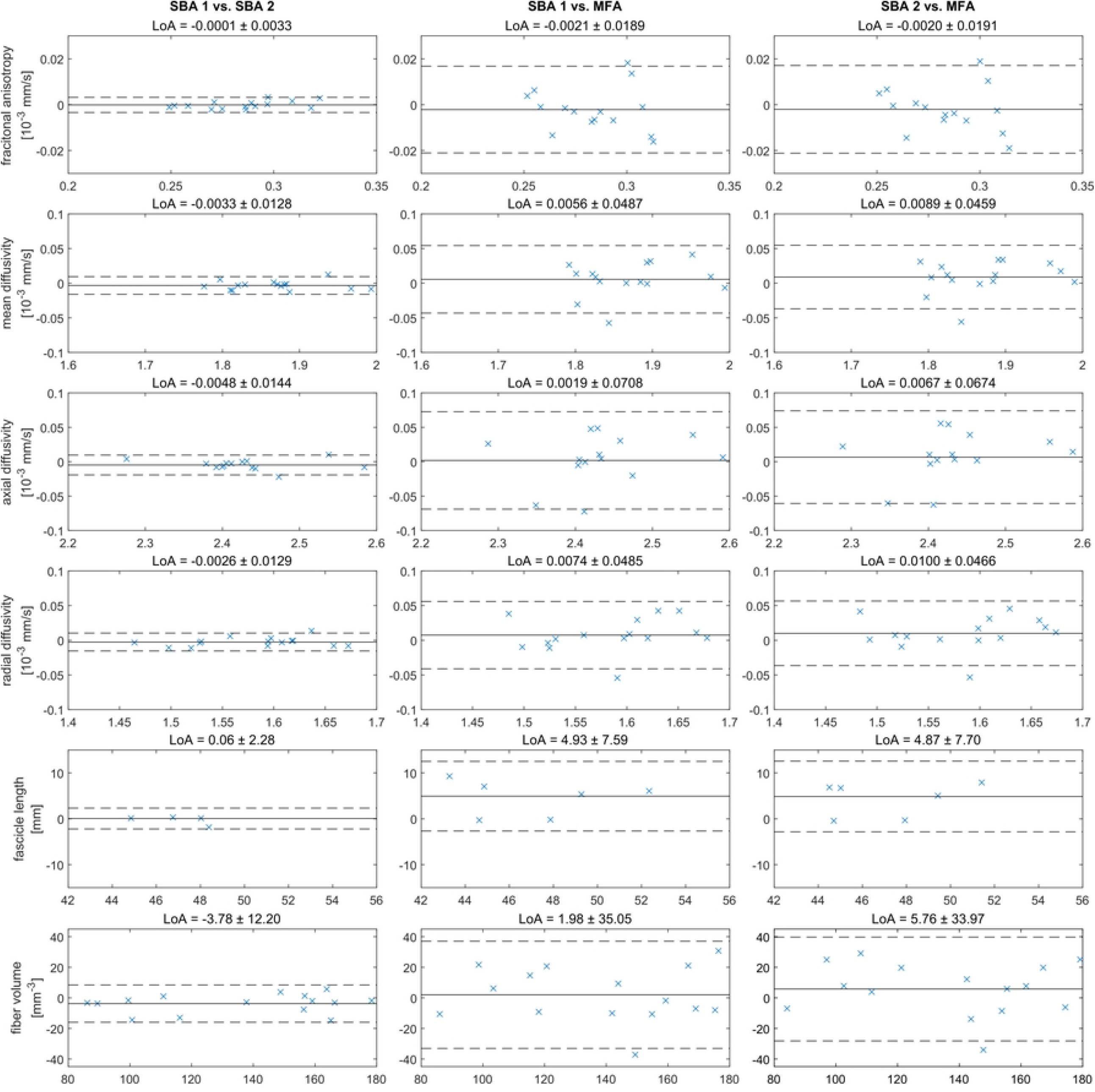
Bland-Altman plots and Limits of Agreement. Bland-Altman plots for interrater-reliability for segmentation-based analysis by two different operators (SBA 1 and SBA 2). and model-free analysis (MFA). The x-coordinate shows the mean of two methods and the y-coordinate shows the value differences between each method. The middle black line shows the mean of the paired differences. The dashed line presents the limits of agreement.

## Discussion

The aim of this study was to investigate the influence of segmentation on muscle tractography. The results show that different segmentation routines and MV results, did not influence SBA fiber tracking with an excellent ICC (≥ 0.975)[44]. This is consistent with Forsting et al.[45] investigating differences between independent raters using pre-specified segmentation routines for the lower limb muscles.. In an explorative comparison of SBA methods to a more convenient model-free fiber tracking (MFA) approach, muscle parameters displayed a good ICC (0.751 – 0.843) and excellent ICC for the mDTI indices (0.901 and 0.93) [44]. Bland-Altman plots (Figure 3) showed no major anomalies in the data and no over- or under-estimation of parameters between SBA and MFA.

Due to the anatomy of the M. supraspinatus and its tendonous structure, differences in MV are to be expected. As the task of both operators was to segment only the muscle and to develop their own segmentation routine independently, they were first introduced to segmentation. The aim was to check if the two operators approached segmentation differently and if this influences a fiber tracking result. As expected, the two operators’ routines differed in the number of processing steps, duration (30-45 minutes), details of manual drawing, and how they defined the muscle-tendon interface (Figure 1). In terms of daily clinical routine, this case may reflect reality, as there is no standardised segmentation routine trained and checked by only one expert[38]. Based on this assumption, ICC estimates were calculated in this study using a two-way random effects model[37,38]. Therefore, the findings reported might be generalised to any random operator performing a muscle segmentation regardless of the operator’s experience and the level of detail in manual segmentation.

The observed differences in FL between SBA and MFA could be attributed to several reasons: Based on the findings of Bolsterlee et al.[28,34] and Damon et al.[26,46] explaining that the most valid tracts are found at the belly of the muscle, we removed tracts that obviously crossed the muscle boundaries in MFA. This may have removed relevant fibers in MFA that were included in SBA. Secondly, MFA used the entire DTI image as the seeding area for fiber tracking, which may result in different density and distribution of tracts[37]. However, this makes it difficult to compare muscle metrics from SBA to MFA and it may be even more interesting to see how accurately these approaches detect changes in muscle architecture over time. Furthermore, cadaveric dissections show that the mean FL for the human M. supraspinatus can vary between 28 mm and 83 mm[39] depending on the region of interest and the number of fascicles measured[19]. This indicates that FL should be compared carefully and in respect to the recruited subjects and the applied analysis methods.

FV is an interesting parameter to interprete the physiological cross-sectional area (PCSA) in combination with MV, FL and the pennation angle of the muscle[19]. As still accepted in science, PCSA seems to be the best predictor of muscle strength[47]. FV representing the PCSA in our case, showed high variance (CV ≥ 21.8) due to very different MV between subjects (68 – 95 mm^3^ in SBA 2). However, the FV was not significantly different between methods (*p* > 0.05) and showed a good ICC (≥ 0.840). Results for FA, MD, AD and RD are comparable to Forsting et al.[37]. Furthermore, our FA and RD values revealed that the recruited subjects did not have irregular values as shown for injured subjects[16,17]. This is important to demonstrate as it appears that muscle tears have a significant effect on FL[12–15] and DTI indices[16,17].

DTI is an innovative opportunity to have a detailed look at a muscle’s fiber architecture. Nevertheless, it has some inherent difficulties. To generate DTI, the diffusion coefficient of water is measured and a mathematical representation of its diffusion is generated in three dimensions. Diffusion should be measured in at least six directions to represent the diffusion anisotropy of the muscle. Fiber tracking therefore maps the connectivity between different points within a biological ROI[46]. However, mDTI acquisition varies between studies in terms of sequences and its settings[46], segmentations[37,38] and data analysis methods[35,36]. This leads to incomparable results between studies[32]. Furthermore, as mDTI measures the water diffusion and allows a reconstruction of tracts, the tractography outcome does not show the actual fiber length[48]. Therefore, a careful plausibility check based on a sum of muscle and DTI metrics is required to allow interpretation of the reconstructed tracts as muscle fibers. While other non-invasive technologies such as conventional 2D ultrasound or micro-computed tomography do not require fiber reconstruction, they appear to be more convenient for daily clinical routine. However, it should be noted that such approaches in interventional studies[49] usually measure a small number of fascicles, which has never been quantitatively tested to represent the muscle architecture accurately[19]. As shown by Charles et al.[19], this may lead to large errors in the interpretation of the whole muscle architecture. Therefore, mDTI appears to be more valid when considering the three-dimensional architecture of the whole muscle. In addition, mDTI can be interpreted as an accurate method for assessing changes after interventions[45].

Comparing mDTI with ultrasound also raises the issue of test economics and feasibility in research and clinical contexts. As mentioned, mDTI involves very complex data processing, which guided the subject of this study. To be precise, usually mDTI contains the following processing steps recommended[34–36]: (1) manual segmentation of target muscle based on anatomical sequences; (2a) resampling and registration of DTI images to T1w images; (2b) quality control of DTI; (2c) definition of ROI-specific stopping criteria for fiber tracking; (2d) integration of the segmented muscle model as ROI; (2e) fiber tracking; (3a) generating a surface model (3b) importing and translating the coordinates of identified tracts onto the surface model; (3c) calculation of muscle parameters. This is a typical workflow in which mDTI appears to be less competitive than ultrasound imaging. Nevertheless, we know that mDTI and ultrasound should be compared carefully[50]. As ultrasound is widely used as a practical imaging tool to detect major irregularities within anatomical structures, mDTI seems to be more important for research tasks due to a high amount of data that can be collected[35]. However, if ultrasound imaging has a higher degree of standardisation in terms of positioning and analysis[21], it may result in a similar level of effort to mDTI. Since a few studies have shown that changes in mDTI parameters can help to detect muscle tears[16,17], we assume that mDTI may be of clinical relevance. The fact that muscle architecture measurements with mDTI have been shown to be reliable[29], and robust[30] gives mDTI advantages over ultrasound imaging[23]. The fact that SBA 1 and SBA 2 did differ in terms of effort and MV but not in terms of fiber tracking outcome, suggests that automatic segmentation approaches[51,52] may be sufficient for accurate calculations of FL.

## Limitations

This study used original Siemens DTI sequences, which have slightly different settings to those recommended for mDTI[40]. This increased the risk of inappropriate mDTI data and incomparable results. In addition, the quality of the data also depends on the size and complexity of the target structure[40]. As the individual anatomy and complexity of the human shoulder complicates DTI acquisition, we observed very heterogeneous data quality, resulting in a large variance between datasets. Furthermore, the analysis of M. supraspinatus also revealed difficulties. As cadaveric dissections have shown, the M. supraspinatus has a very different fiber orientation within the anterior or posterior superficial, middle and deep regions[39]. Since it is known that a tractography can reveal disoriented tracts and tracts that cross the muscle boundaries[30], we were not able to detect in detail the different fiber orientation within the target muscle. Furthermore, as our commercial sequences deliberately showed untouched settings to test feasibility in a clinical context, this led to difficulties in resampling and registering the DTI images to the T1w images used for segmentation. We therefore decided not to calculate other muscle parameters such as pennation angle, which has also been discussed elsewhere[35]. The fact that our study results are based on a small sample size including highly active student athletes means that the results should be interpreted with caution alongside the other studies mentioned[34,35].

## Conclusion

This study shows that different segmentation routines do not influence tractography for the human shoulder M. supraspinatus. This is in concordance to a study for the lower limb[37]. Furthermore, tractography without a segmentation step (MFA) shows good ICC estimates. Therefore, SBA or MFA may be advantageous depending on the needs. In the case of a muscle lesion[16,17], MFA can be used as a first and quick decision basis for further treatment. On the other hand, SBA can be used to show training-induced changes and for a more precise analysis of muscle parameters. In terms of test economy, MFA may compete with ultrasound imaging methods. While SBA 1 and SBA 2 differed in time and detail, but not in tractography results, automatic segmentation approaches can be considered to maintain the accuracy of FL calculations and speed up analysis. As the human shoulder joint is known to be a challenge for any imaging technique, the results of mDTI were very promising and detailed. Therefore, mDTI can claim practical relevance. Furthermore, mDTI seems worth considering as a standard sequence within a musculoskeletal MR application in a clinical context. The evaluated mDTI processing methods could form the basis for further interventional studies and future methods. MFA is an acceptable and reliable mDTI analysis method that is reduced to a minimum number of processing steps to assess the muscle fiber architecture and quality as quickly as currently possible. To our knowledge, this is the first study using mDTI in the human shoulder in this context. It remains unclear whether MFA could be applied to other muscles with different fiber orientations.

## Acknowledgments

We thank all the radiologists and medical scientists involved in this study. We also thank Frank Yeh for software support from DSI Studio.

## Data availability

The data that support the findings may be obtained from the corresponding author.

